# Unexpected gene activation following CRISPR-Cas9-mediated genome editing

**DOI:** 10.1101/2021.06.14.448328

**Authors:** Anna G Manjón, Hans Teunissen, Elzo de Wit, René H Medema

**Affiliations:** The Netherlands Cancer Institute; Division of Cell Biology; Division of Gene Regulation

## Abstract

The discovery of the Clustered Regularly-Interspaced Short Palindromic Repeats (CRISPR) and its development as a genome editing tool has revolutionized the field of molecular biology. In the DNA damage field, CRISPR has brought an alternative to induce endogenous double-strand breaks (DSB) at desired genomic locations and study the DNA damage response and its consequences. Many systems for sgRNA delivery have been reported in order to efficiency generate this DSB, including lentiviral vectors. However, some of the consequences of these systems are yet not well understood. Here we report that lentiviral-based sgRNA vectors can integrate into the endogenous genomic target location, leading to undesired activation of the target gene. By generating a DSB in the regulatory region of the *ABCB1* gene using a lentiviral sgRNA vector, we can induce the formation of taxol-resistant colonies. We show that these colonies upregulated *ABCB1* via integration of the *EEF1A1* and the U6 promoters from the sgRNA vector. We believe that this is an unreported CRISPR/Cas9 artefact that researchers need to be aware of when using lentiviral vectors for genome editing.

## Introduction

The discovery of the Clustered Regularly-Interspaced Short Palindromic Repeats (CRISPR), their role in the prokaryotic immune system and subsequent development as a genome editing tool has revolutionized the field of molecular biology^1–6^. In recent years, many laboratories have developed CRISPR/Cas9 as a tool that can be applied to study many different biological questions^7^. In the DNA damage field, CRISPR has brought an alternative to induce endogenous double-strand breaks (DSB) at desired genomic locations. This system allowed for the study of the DNA damage response and its consequences in different genome compartments or structures^8^. Combining imaging and high throughput technologies with DSB-induced Cas9 systems allows one to examine processes such as transcription, chromatin dynamics, and DNA replication.

The CRISPR/Cas9 system needs to be delivered in an accurate manner for efficient gene editing. On the one hand, the Cas9 protein needs to be expressed in the host system or delivered in a form of a Ribonucleoprotein (RNP) complex^9^. On the other hand, a target-specific single guide RNA (sgRNA) – formed by CRISPR RNA (crRNA) and transactivating CRISPR RNA – needs to direct Cas9 to the target site^10^. It is important to choose the right delivery strategy for the sgRNA to survive the degradation processes in the cell and translocate into the nucleus to allow for gene editing. To date, we can classify sgRNA delivery methods into viral and non-viral, based on whether viral constructs are used for transfection^7^.

Viral vectors include adeno-associated viruses (AAVs) and lentiviruses (LVs). Specially in LVs, Cas9 and sgRNA are relatively easy to clone, produce and efficiently transduced into the host cell. However, the bigger challenge of these systems is the random integration of the construct into the genome^11^. We can divide the non-viral methods into physical and chemical. Physical methods include microinjections – where the sgRNAs are directly injected by a needle – and electroporation – where electric currents open the cell membrane for the delivery of molecules into the cell^12,13^. Chemical delivery methods comprise a DNA or mRNA form of the sgRNA that can be transfected into the host by lipofectamine reagents^14^. In these latter strategies the transfection efficiency can be lower, but they are a safer alternative, as random viral integrations do not occur.

Even though targeting genomic regions with the CRISPR/Cas9 system is tightly controlled and specific, it is known that off-target cutting activity could still occur^5,15,16^. Other limitations of CRISPR include the requirement for a protospacer adjacent motif (PAM) to the target DNA sequence and the DNA-damage toxicity triggered after the CRISPR-induced DSB^17^. Nonetheless, valuable efforts have been made to understand and minimize these drawbacks. However, much less is known about how viral CRISPR/Cas9 delivery methods may affect genome integrity and gene expression when randomly integrated into the host genome.

Here we show that a LV-based sgRNA vector can integrate into the endogenous genomic target location thereby affecting gene expression of the target gene. By generating a DSB in the regulatory region of the *ABCB1* gene with this system, we can produce taxol resistant clones that upregulated *ABCB1* through transcriptional activation via the *EEF1A1* and the U6 promoters from the sgRNA vector. We believe that this unreported gene activation mechanism following CRISPR-Cas9-mediated genome editing needs to be taken into consideration when inducing DSBs with a sgRNA lentiviral method.

## Results

### A LentiGuide-induced DSB in the ABCB1 promoter leads to upregulation of ABCB1

We have previously shown that in RPE-1 cells, the major mechanism of taxol resistance is transcriptional activation of the *ABCB1* gene, that encodes for the multi-drug resistance protein MDR1 or P-Glycoprotein (PgP)^18,19^. Using the lentiviral system lentiGuide-Puro from the Zhang Lab, we cloned different sgRNAs targeting different non-coding regions across the *ABCB1* locus to induce a DSB (**Fig 1A**). We chose non-coding regions to avoid the possibility that a break-induced change in coding sequence could result in acquired taxol resistance. 7 days after lentiviral infection and puromycin selection we treated the RPE-1 cells with 8nM of taxol in order to select cells that over-expressed PgP. Surprisingly, we observed that only cells treated with sgRNAs targeting the promotor of *ABCB1* became resistant to taxol (**Fig 1A**), as we observed a considerable number of RPE-1 colonies growing under taxol pressure. In order to better understand the mechanisms responsible for the acquisition of the taxol-resistant phenotype, we decided to individually characterize the taxol-resistant clones from the sgRNA targeting the *ABCB1* promoter. Therefore, we expanded under taxol pressure the resistant colonies observed in the colony outgrowth assays. When performing a viability assay with increasing doses of taxol, we observed that all clones were resistant to high concentrations of taxol, and could be re-sensitized with Tariquidar, a PgP inhibitor (**Fig 1B**). As expected, with Western Blot and qRT-PCR assays we could confirm that the taxol-resistant clones expressed high levels of PgP as well as mRNA respectively (**Fig 1C-D**). Thus, confirming that the mechanism of taxol resistance was through *ABCB1* upregulation. By performing intronic smRNA-FISH, which allows for visualization of active transcription sites, we demonstrated that only one allele was actively transcribing *ABCB1* (**Fig 1E**), confirming that *ABCB1* copy number amplifications were not observed in these clones.

**Figure 1.**
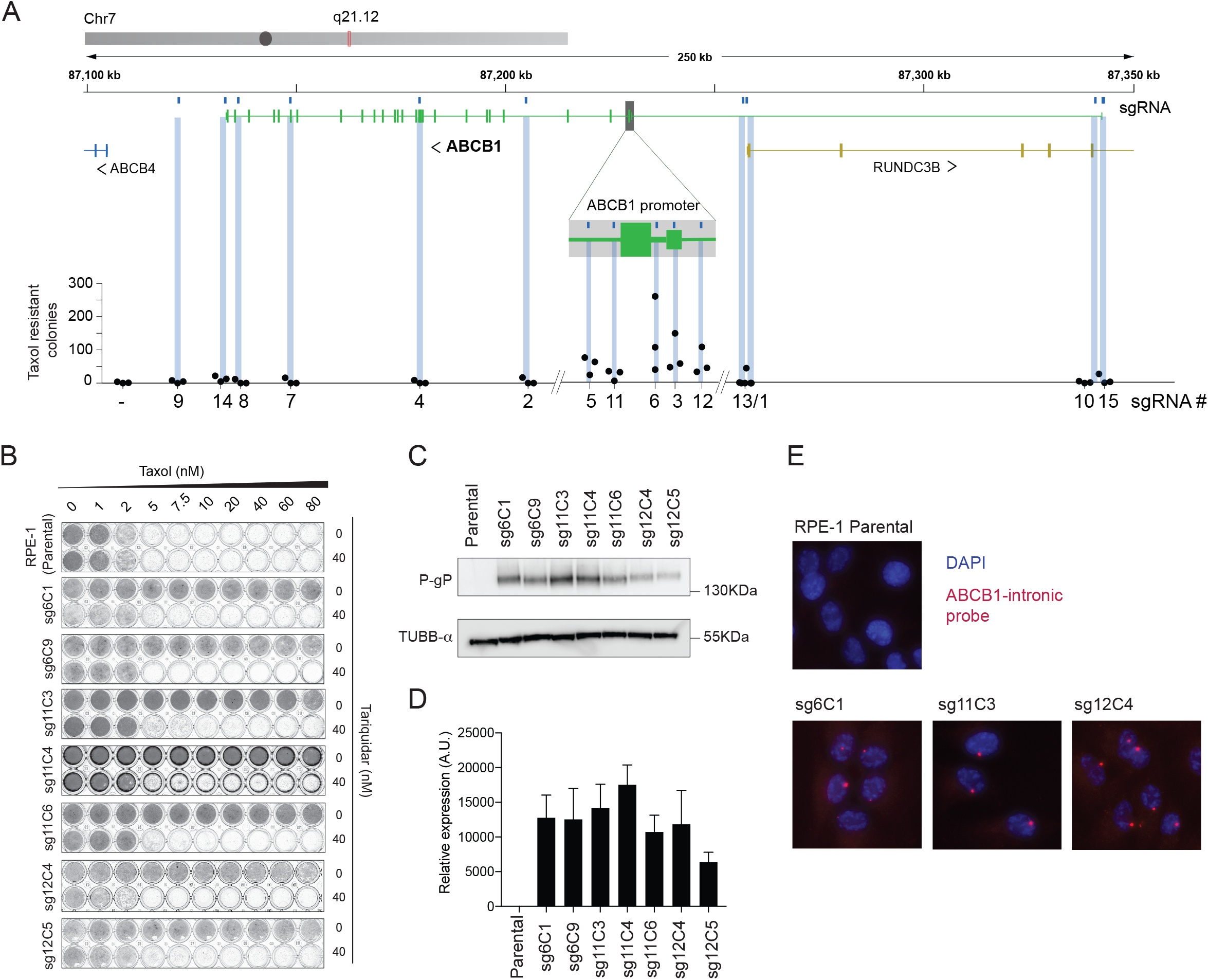
**A)** Graphical representation of the ABCB1 genomic region and the location of the gRNA targeting the gene. RPE-1 cells were infected with a Lentivirus carrying one of the gRNAs and after 7 days of puromycin selection 1 milion cells were plate with 8nM of Taxol for Colony Outgrowth Assay. Taxol resistant cells were counted an plotted in the graph. N=3. **B)** Crystal violet staining of viability assay on RPE-1 Parental cells and taxol resistant clones obtained from A. For the clones’ nomenclature, sg# represents the sgRNA from where they are derived and C# the clone number. **C)** Western Blot showing the levels of the PgP and control (α-TUBB) upon in RPE-1 Parental and taxol resistant clones. **D)***ABCB1* mRNA levels determined by qRT-PCR and normalized to *GAPDH* expression levels, n=2. Error bars show the SD. **E)** Representative smRNA-FISH images of RPE-1 Parental and clones for the *ABCB1* gene and DAPI. The images are projections of 0,5μm sections and a total 5μm in thickness. Scale bar, 15μm.

### The LentiGuide vector integrates and drives gene expression upon a DSB in the ABCB1 promoter

To exclude that DNA translocations or insertions might be induced by the DSB and could modify the activity of the *ABCB1* promoter, we performed Targeted Locus Amplification (TLA), a chromosome conformation capture-based technique, enabling the identification of single nucleotide variation and genomic rearrangements in a specific locus using a single PCR reaction^20^. We selectively amplified and sequenced the DNA flanking the *ABCB1* promoter. We compared RPE-1 Parental cells with a taxol-resistant clone derived from the sgRNA #6 targeting the promoter of *ABCB1* (sg6C9). Surprisingly, we found that our TLA experiments for the *ABCB1* promoter amplified a 1.3kb region from chromosome 6 in the taxol-resistant clone (**Fig 2A**, green arrow). When zooming in on that region, we discovered that the promoter of the *EEF1A1* gene was amplified in the sg6C9 taxol-resistant clone (**Fig 2B**). The read distribution over the *EEF1A1* promoter is reminiscent of genomic insertions previously mapped with TLA^20^. To confirm the fusion of the *ABCB1* and *EEF1A1*, we performed PCRs on genomic DNA using either *Forward* and *Reverse* primers amplifying the *ABCB1* break site or a *Forward* primer binding the promoter region of *EEF1A1* together with a *Reverse* from the *ABCB1* promoter. Only when *EEF1A1* and *ABCB1* are juxtaposed in the genome this will result in a PCR product (**Fig 2C**). Remarkably, we found out that not only the taxol-resistant clone sg6C9 but also all the other clones derived from the sgRNA #6 and some others from #3, #5, #11 and #12, all generating a DSB in the promoter of *ABCB1*, gave a PCR product when using the *ABCB1* and *EEF1A1* primers (**Fig 2D**). We could also observe a higher band appearing when amplifying the sequence over the break site with *Forward* and *Reverse ABCB1* primers (**Fig 2E**). These data confirm that the *EEF1A1* promoter was integrated in the break site in the regulatory region of *ABCB1*. When we sequenced the PCR products from the different clones, we observed that there were other sequences belonging to the U6 promoter and the puromycin-resistant cassette integrated (data not shown). We next decided to align the sequence reads of the TLA experiment analyzing the sg6C9 taxol-resistant clone to the LentiGuide vector sequence that was used to clone the *ABCB1*-targeting sgRNAs to induce the DSB. We found that in the sg6C9 taxol-resistant clone, there was a large region aligning with the LentiGuide vector, suggesting that the *EEF1A1* integration found in the *ABCB1* promoter belonged to the LentiGuide vector and not to the gene found on chromosome 6 (**Fig 2F**). We therefore conclude that the lentiGuide-Puro vector had been integrated into the *ABCB1* promoter, most likely due to the presence of the CRISPR-induced DSB in that region. As the U6 promoter is a RNA Pol III promoter most likely this will not result in mRNA and protein translation. Therefore, most probably the *EEF1A1* promoter from this vector induced the transcriptional activation of *ABCB1.*

**Figure 2.**
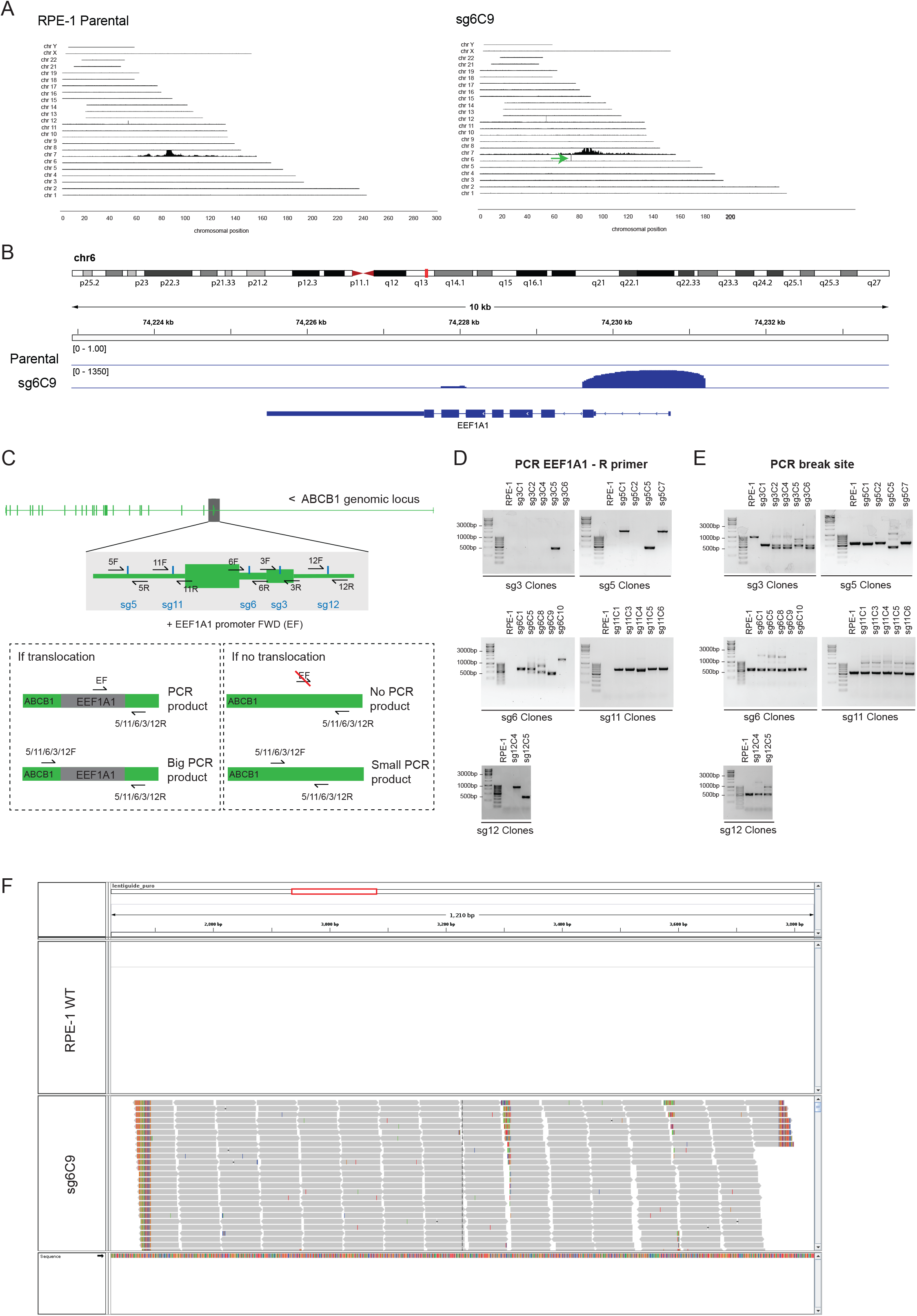
**A)** TLA analysis for *ABCB1* contacts in RPE-1 Parental and taxol resistant clone AB6C9 covering the whole genome. Green arrow in AB6C9 shows a *de novo* interaction found between *ABCB1* and a region in chromosome 6. **B)** TLA analysis for RPE-1 Parental and AB6C9. Zoom in in the region of chr6 with de novo interaction for AB6C9. **C)** Graphical representation of the gRNA targeting the *ABCB1* promotor region. A common FWD primer binding EEF1A1 promoter was used in combination with different REV primers for each of the gRNA break sites. Only when EEF1A1 is integrated in cis we will obtain a PCR product. **D)** PCR products using the primers in C over the *ABCB1* and *EEF1A1* regions in RPE-1 Parental and the different taxol resistant clones. **E)** IGV screen shot where the reads of the TLA experiments are aligned to the lenti-guide puro sequence. In AB6C9, reads align to the lenti-guide vector.

## Discussion

We show here that a lentiviral sgRNA delivery system to induce a DSB close the transcriptional start site of a gene can result in integration of the vector in the break site, and activation of the gene. When generating a DBS in the regulatory region of *ABCB1* with this system, we were able to find cells with genetic alterations that contained the U6 and *EE1A1* promoters of the lentiviral vector. We believe that the DSB increased the probability of the vector to integrate into this location. In RPE-1 cells *ABCB1* is repressed and the cells are sensitive to taxol. The integration of these promoters allowed for gene activation and produced a taxol resistant phenotype (**Figure 3**). This mechanism does not appear to be of high frequency, but selection of cells with high transcriptional levels of *ABCB1* by taxol increased its occurrence. Nonetheless, as seen by colony formation outgrowth, we found that this event may happen in up to three cells out of a thousand, suggesting that this type of genetic alterations have to be taken into consideration.

**Figure 3.**
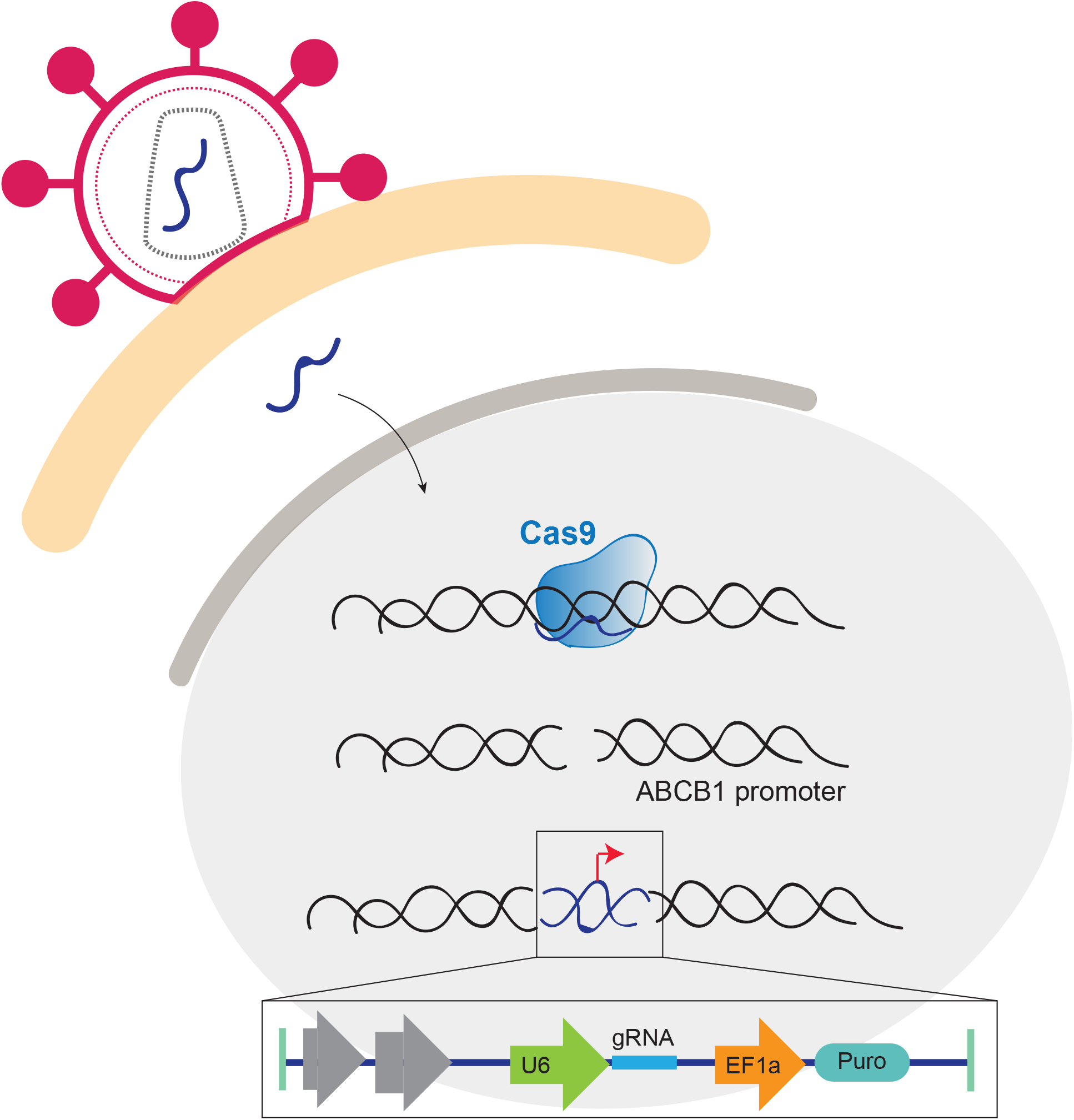
When a lenti-guide Puro vector is used to deliver a gRNA to induce a DSB, the gRNA can be integrated into the break site. The DSB was induced in the promoter of ABCB1 and therefore the highly active promoters of the vector were driving the expression of the ABCB1 gene. In this case, we were selecting for cells that upregulated ABCB1 and therefore the frequency of this event was higher.

HIV-1-based lentiviral vectors convert single-strand RNA into double-strand DNA by reverse transcription and subsequently insertion into the genome of post-mitotic cells^7^. Lentiviral vectors have become important tools to deliver components of the CRISPR/Cas9 system for genome editing. However, in gene therapy, stable viral integrations come with concerns regarding safety^21^. Among them, the deregulation of genes caused by the insertions and mutagenesis found in gene therapy for immunodeficiencies in patients^22^.

Many researchers are currently using lentiviral vectors for delivery of CRISPR/Cas9 components, as sgRNAs are relatively easy to clone into them^7^. Lentiviral sgRNA-delivery systems are used in functional genetic screens to find lethal interactions of specific biological processes^23^. Even though many limitations are known regarding off targets or difference in efficiency between sgRNAs^24–26^, little is known about how lentiviral-based CRISPR can affect gene transcription changes. We speculate that targeted viral integration could result in deregulation of genes that may affect biological functions and therefore lead to false positive candidates when performing functional screens. Thus, when performing screens, it is important to have a good sgRNA complexity and reproducible results.

Furthermore, as CRISPR enables the induction of DNA breaks at specific endogenous loci, more and more researchers are using several Cas9 systems to study of DSB repair and its biological consequences^8,27,28^. As we show here, inducing a DSB in a gene regulatory region could have consequence in gene expression thus leading to incorrect interpretation of the results. Therefore, to study long term consequence of the DNA damage response we suggest to employ non-integrative systems such as synthetic gRNAs delivered in an RNA form^29^.

## Materials and Methods

### Cell lines and cell culture conditions

hTert-immortalized retinal pigment epithelium (RPE-1) and derived cell lines were maintained in DMEM/F-12 + Glutamax (Gibco, Life Technology) supplemented with 1% penicillin/streptomycin and 6% fetal bovine serum (FBS, S-FBS-EU-015, Serana).

### Taxol and Tariquidar treatment

Taxol and Tariquidar were dissolved in DMSO and prepared at stock concentrations before usage at varying final concentrations as indicated in each figure.

### sgRNA designed and cloning

the gRNAs targeting *ABCB1* were cloned into a lenti-guidePuro (Addgene plasmid # 52963) using the BsmBI restriction site. sgRNA sequences are summarized in **Table 1**.

**Table 1.**
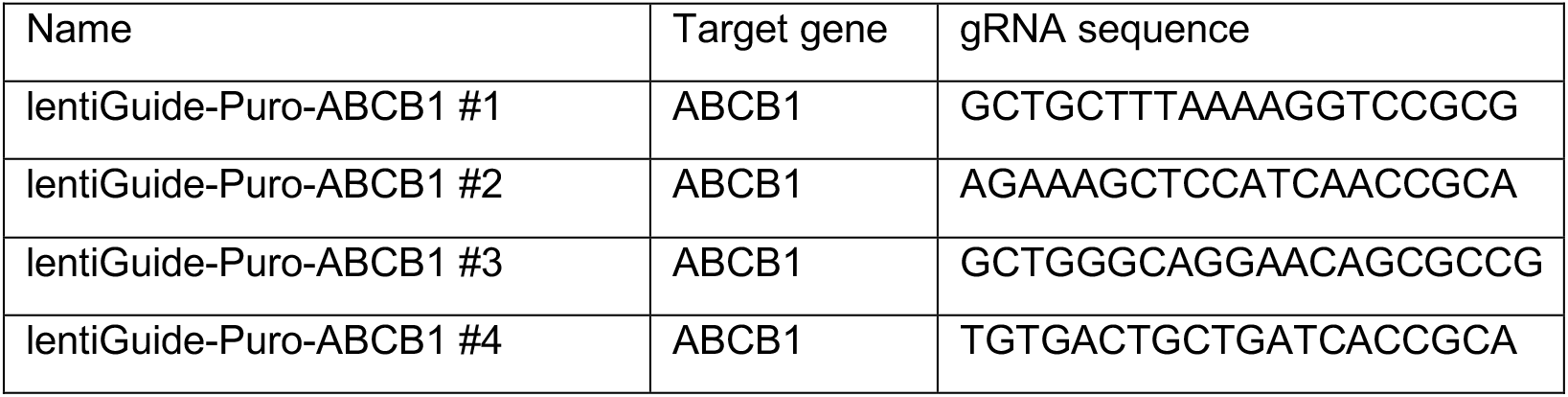
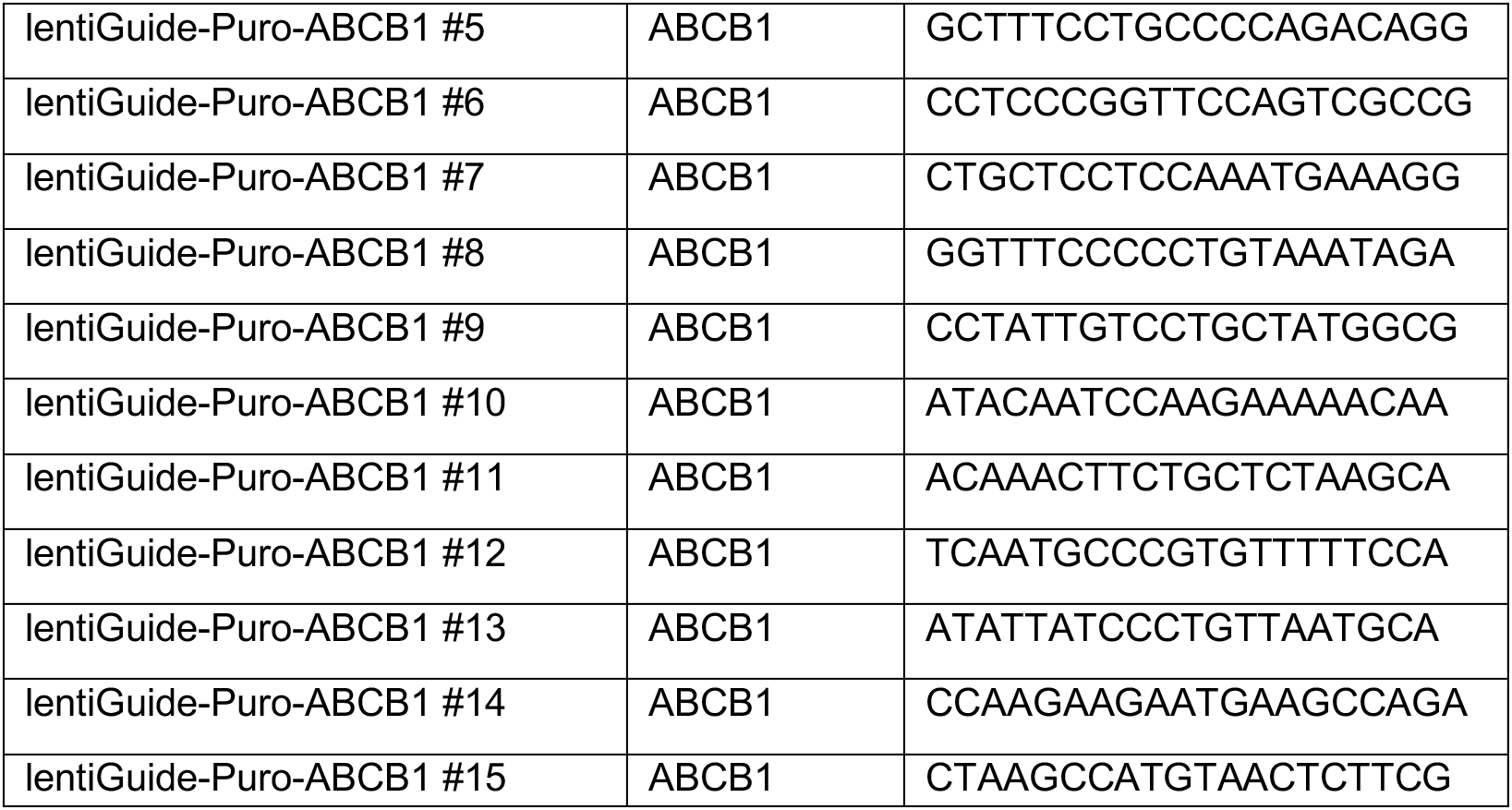
gRNA sequences targeting ABCB1

### Colony Outgrowth Assays

1 million cells were treated with 8nM of taxol and allowed to grow out for 15 days. Plates were fixed in 80% Methanol and stained with 0.2% Crystal Violet solution. After fixation, the number of taxol resistant cells were counted.

### Viability assays

For viability assays, 1000 cells were plated in a 96-well plate and treated for 7 days with indicated drug concentrations. Subsequently, plates were fixed in 80% Methanol and stained with 0.2% Crystal Violet solution.

### RNA isolation and qRT-PCR analysis

RNA isolation was performed by using Qiagen RNeasy kit and quantified using NanoDrop (Thermo Fisher Scientific). cDNA was synthesized using Bioscript reverse transcriptase (Bioline), Random Primers (Thermo Fisher), and 1000 ng of total RNA according to the manufacturer’s protocol. Primers were designed with a melting temperature close to 60 degrees to generate 90–120-bp amplicons, mostly spanning introns. cDNA was amplified for 40 cycles on a cycler (model CFX96; Bio-Rad Laboratories) using SYBR Green PCR Master Mix (Applied Biosystems). Target cDNA levels were analyzed by the comparative cycle (Ct) method and values were normalized against GAPDH expression levels. qRT-PCR oligo sequences are summarized in **Table 2**.

**Table 2.**
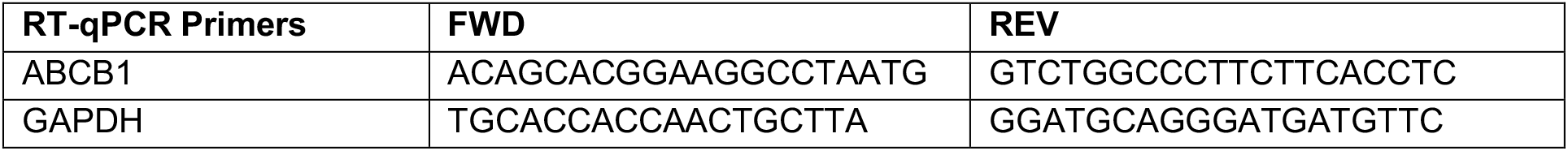
RT-qPCR primers

### Western Blots

For western blot experiments, equal amounts of cells were lysed with Laemmli buffer and separated by SDS–polyacrylamide gel electrophoresis followed by transfer to a nitrocellulose membrane. Membranes were blocked in 5% milk in PBST for 1h at RT before overnight incubation with primary antibody in PBST with 5% BSA at 4°C. Membranes were washed three times with PBST followed by incubation with secondary antibody in PBST with 5% milk for 2h at RT. Antibodies were visualized using enhanced chemiluminescence (ECL) (GE Healthcare). The following antibodies were used for western blot experiments: α-Tubulin (Sigma t5168), MDR(PgP) (sc-8313). For secondary antibodies, peroxidase-conjugated goat anti-rabbit (P448 DAKO, 1:2000), goat anti-mouse (P447 DAKO, 1:2000) and rabbit anti-goat (P449) were used.

### smRNA FISH

RPE-1 cells were plated on glass coverslips and washed twice with BS before fixation in 4% PFA in PBS for 10 minutes at room temperature. After two additional washes in 1x PBS coverslips were incubated in 70% ethanol at 4°C overnight. Coverslips were incubated for pre-hybridization in wash buffer (2x saline-sodium citrate (SSC) with deionized formamide (Sigma) 10%) for 2-5 minutes at room temperature. RNA FISH probe mix wash dissolved in hybridization buffer (wash buffer supplemented with 10% dextran sulfate). 38 probes labelled with Cy5 were targeted to the intronic regions of ABCB1 (Biosearch technologies). Coverslips were incubated in hybridization solution for at least 4h at 37°C. Then coverslips were washed twice for 30 minutes with wash buffer followed by a quick rinse with 2x SSC. Finally, coverslips were washed once for 5 minutes in 1x PBS before mounting on slides using Prolong gold DAPI mounting medium (Life Technologies). Images were acquired with the use of a DeltaVision Elite (Applied Precision) equipped with a 60x 1.45 numerical aperture (NA) lens (Olympus) and cooled CoolSnap CCD camera. ABCB1 transcription start site quantification was performed manually double blind.

### TLA analysis

TLA was performed as previously described with minor modifications. TLA libraries were sequenced on a MiSeq and were mapped to genome using bwa bwasw[Heng Li ref: PMID: 20080505] to enable partial mapping of sequence reads. Reads were mapped to hg19 reference of the human genome.

